# Detection, Persistence, and Antimicrobial Resistance Profiles of *Escherichia coli* and Shiga Toxin-Producing *E. coli* in Nut Orchards with Integrated Grazing Animals

**DOI:** 10.64898/2026.11.13.766593

**Authors:** Joanna G. Rothwell, Sejin Cheong, Cory L. Schlesener, Craig C. Miramontes, Claire Shaw, Richard V. Pereira, Bart C. Weimer, Alda F. A. Pires

**Affiliations:** Department of Population Health and Reproduction; 100K Pathogen Genome Project, School of Veterinary Medicine, University of California Davis, Davis, CA 95616, USA

**Author notes:** Corresponding author: Alda F. A. Pires^1^.

**Keywords:** Integrated crop-livestock systems, grazing, soil, sheep, nut orchards, Shiga toxin-producing *Escherichia coli*, whole genome sequencing, antimicrobial resistance, foodborne pathogens

## Abstract

Integrating grazing animals into nut orchards can introduce foodborne pathogens including *Salmonella enterica* and Shiga toxin-producing *Escherichia coli* (STEC) into crop production environments. This study investigated pathogen transfer and persistence in the soil following livestock grazing (sheep/chicken) in two organic almond orchards and one walnut orchard in California. Soil and fecal samples were collected periodically for 120 days after grazing. Enrichment, selective culture and PCR were used for STEC and *Salmonella enterica* detection. Antimicrobial resistance (AMR) was assessed using microdilution assays, and whole genome sequencing (WGS) was used to characterize AMR genes, virulence genes and population structure. All 284 soil and 79 fecal samples collected tested negative for *Salmonella enterica* and *E. coli* O157. Presumptive STEC was detected by colony PCR in 3.2% (n = 9/284, 95% CI 1.7-5.9%) of the soil samples, and 34.2% (n = 27/79, 95% CI 24.7-45.2%) of fecal samples. Shiga toxin genes were confirmed by WGS in 15 of 31 sequenced isolates. Predominant subtypes were *stx*_1c_ and *stx*_2b_, and O176:H4 and O174:H2 were the most common serotypes. Phenotypic AMR was detected in 33.3% (n = 12/36) of isolates, with multidrug resistance in 8.3% (n = 3/36). A soil and a sheep fecal isolate differed by one core-genome single nucleotide polymorphism and shared a closed multidrug-resistance plasmid of identical gene content, consistent with a lineage resident in the flock that transferred to soil. Confirmed STEC in orchard soil was rare and transient. These results provide evidence to inform food safety guidelines on crops integrated with livestock.

**Importance:** Farmers use grazing animals including chicken and sheep in nut orchards to improve the health of the soil and productivity of the orchard while also increasing sustainability. However, animals can shed bacteria that cause foodborne disease, and as nuts come into contact with the orchard soil before harvest, bacteria surviving in the orchard could contaminate the crop. Growers wait a fixed period between grazing and harvest, however bacterial survival may vary with climate, soil conditions and animal species. In this study, Shiga toxin-producing *E. coli* was common in sheep feces but was rarely confirmed in orchard soil, and most bacteria isolated from chicken feces did not carry Shiga toxin genes when their genomes were sequenced. The bacteria of greatest concern, *Salmonella enterica* and *E. coli* O157, were not found in any samples. This research can inform grazing guidance for nut orchards, supporting both food safety and sustainable farming.

## Introduction

Integrated crop-livestock systems (ICLS) combine grazing animals with crop production environments for synergistic animal-crop benefits and enhanced farm sustainability (1,2). In these systems, animals consume cover crops or volunteer crops, reducing the costs associated with mowing and animal feed. Additionally, animals produce manure that is deposited directly into the soil, increasing nutrient cycling and soil quality while also reducing the need for fertilizers (3–7). Fruit and nut orchards are particularly suitable for incorporation with grazing animals as widely spaced trees allow sufficient light for productive forage or cover crop growth (8). Additionally, grazing animals reduce pest pressure through consuming residual fruits and nuts that may be reservoirs of pests and plant disease (9,10). Economically, ICLS benefit farms through diversification of income sources, increasing the use of a single piece of land, and align with profitable organic production requirements. Based on these advantages, there is growing interest in ICLS in orchards as a form of regenerative agriculture (11).

However, the integration of livestock into food production environments has important food safety considerations. Raw manure produced by animals can contain pathogens associated with foodborne disease such as Shiga toxin-producing *Escherichia coli* (STEC), *E. coli* O157:H7, and *Salmonella enterica*. In particular, small ruminants and chicken are known to be asymptomatic reservoirs for STEC and *Salmonella enterica,* respectively, and these animals intermittently shed bacteria into the environment (12,13). Tree nuts may come into direct contact with the soil during harvest making soil an important medium for potential cross-contamination. A recent outbreak of *E. coli* O157:H7 was linked to walnuts in 2024, causing 13 illnesses across two states including two cases of hemolytic uremic syndrome. This was the first documented outbreak linked to walnuts in the United States, and the route of contamination was not established despite product and environmental sampling at the implicated grower and processor (14). There have also been several outbreaks of *Salmonella* linked to almonds in the United States and internationally (15–17). Almonds grown in California and sold in North America since 2007 are required to be processed with a treatment achieving a 4-log reduction of *Salmonella* (18) to reduce the risk of outbreaks, although the same requirements are not enforced in other nut types or for small growers selling almonds directly to consumers. Pathogens may survive in soil at low levels for extended periods (19–23), and can be transferred to the crop when nuts are dropped and dried on the orchard floor prior to harvest (24). To help reduce the likelihood of contamination of crops for human consumption, many growers follow the National Organic Program guidelines from the United States Department of Agriculture (USDA) of a 120 or 90-day waiting period from raw manure application to harvesting (120 days if in contact with soil or 90 days if crops do not contact soil) (25). However, specific rules surrounding grazing systems are not currently established. Under the FSMA Produce Safety Rule, FDA withdrew the proposed nine-month interval for untreated biological soil amendments of animal origin and currently imposes no mandatory interval, stating only that it does not object to growers following the National Organic Program standards (26). Manure deposited *in situ* by grazing animals falls instead under the animal-intrusion provisions, which require growers to assess growing areas and not harvest produce likely to be contaminated (27). Neither approach is based on data from tree nut orchards. Therefore, more research is needed to quantify pathogen transfer and survival over time in orchard soil from grazing to harvest.

The aims of this study were to: 1) characterize the prevalence of foodborne pathogens in grazing animal feces and nut orchard soils after livestock (sheep and chicken) grazing; 2) determine if these pathogens are persisting in this environment; and 3) identify the antimicrobial resistance and virulence profiles of isolated pathogens. This information will be used to develop strategies for mitigating these risks while promoting sustainable farming practices and better understanding the interactions between agricultural practices, bacterial pathogens, and animal and human health.

## Results

### Prevalence of pathogens in soil and fecal samples

No *Salmonella enterica* or *E*. *coli* O157 was isolated from fecal or soil samples collected from the organic almond and walnut farms during the 2021-2022 pilot study. Across the two-year study period, 9 of the 284 soil samples (3.2%, 95% CI 1.7-5.9%) and 27 of the 79 fecal samples (34.2%, 95% CI 24.7-45.2%) were positive for non-O157 STEC. The prevalence observed at each field and sampling timepoint is summarized in Table 1. Eight of the nine PCR-positive soil samples were sequenced, and Shiga toxin genes were confirmed in one of these, giving a WGS-confirmed STEC prevalence in soil of 1 in 284 samples (0.4%, 95% CI 0.1-2.0%), with one PCR-positive soil sample not sequenced. Of the 27 PCR-positive fecal samples, 23 were sequenced and passed QC and Shiga toxin genes were confirmed in 14, giving a WGS-confirmed fecal STEC prevalence of 14 in 79 samples (17.7%, 95% CI 10.9-27.6%).

**Table 1:**
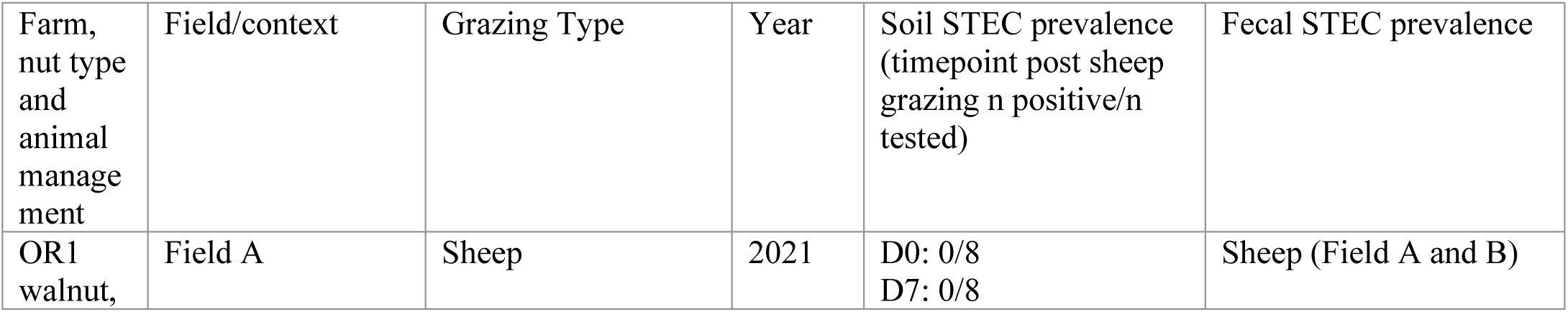
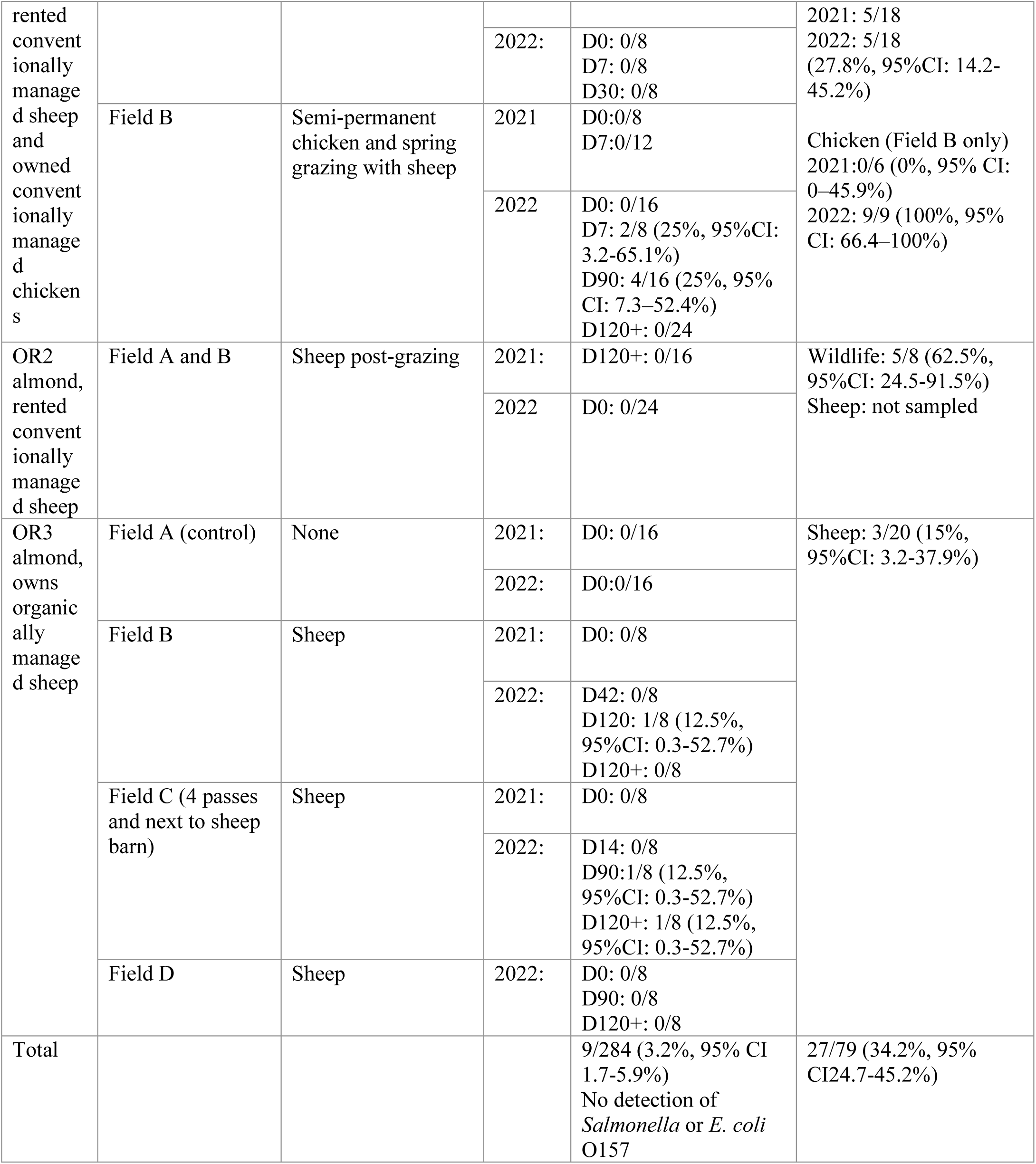
Prevalence of pathogens in soil and fecal samples collected from three organic nut orchards over 2021 and 2022. Soil sampling time points are given as days post-grazing (D0 = immediately post-grazing). No *Salmonella enterica* or *E. coli* O157 was detected in any sample. Prevalence is based on colony PCR and is presumptive; Shiga toxin genes were confirmed by whole genome sequencing in a subset of these isolates, as reported in the Results. Wilson 95% confidence intervals were used for overall prevalence and Clopper-Pearson exact 95% confidence interval for field-timepoint prevalence. Upper 95% confidence limits for zero positive samples were 36.9% (0/8), 26.5% (0/12), 20.6% (0/16), and 14.2% (0/24).

No soil contamination of OR1 Field A was observed despite STEC detection by PCR in 27.8% of sheep fecal samples from this farm. STEC was detected in 4.8% (n = 6/124) of total soil samples from OR1, all from Field B where chicken tractors were continuously present. These STEC were isolated only in 2022 at D7 (n = 2/8, 25%) and D90 (n = 4/16, 25%) post-sheep grazing and were no longer detected by harvest. At OR2, where sheep had grazed the fields 120 days prior, pathogens were not detected in soil samples or during sampling a year after the first grazing event. Wildlife (deer) feces collected during the first visit of OR2 at harvest time were 62.5% (n = 5/8) positive for STEC. STEC was detected in 2.5% (n = 3/120) of the soil samples collected from OR3 with sporadic detections in Field B at D120 and Field C at D90 and D120+. Field C is directly adjacent to permanent sheep housing, and sheep escaped into this field before the D120+ sampling.

### Phenotypic antimicrobial resistance distributions

The phenotypic antimicrobial susceptibility of STEC isolates was tested against the 14 antimicrobials on the CMV3AGNF panel, of which 13 could be interpreted, using broth microdilution assays with results summarized in Table 2 (full data listed in Supplementary Table S1). Of the 36 isolates tested, 33.3% (n = 12/36) were resistant to one or more antimicrobial, with resistance observed across all sample types. Resistance was most prevalent in chicken isolates (n = 4/9, 44.4%), followed by deer (n = 2/5, 40%), sheep (n = 4/13, 30.8%) and soil (n = 2/9, 22.2%). OR1 had the highest AMR prevalence (n = 10/25, 40%) while OR2 was limited to two deer isolates with cefoxitin and streptomycin resistance, one of which was also resistant to ceftriaxone and ceftiofur, and there was no AMR observed in OR3 isolates. Three isolates (8.3%) demonstrated multidrug resistance (resistance to ≥3 antimicrobial classes) all originating from OR1, including matching chloramphenicol, tetracycline and streptomycin profiles in one sheep and one soil isolate, and the most complex resistance profile of cefoxitin, tetracycline, ceftriaxone and streptomycin in a single chicken isolate. Phenotypic resistance to cefoxitin was observed in sheep (n = 1), deer (n = 2), and chicken (n = 2) isolates but not in soil. Gentamicin resistance was observed in 4 of 36 isolates (11.1%), from sheep (n = 2), chicken (n = 1) and soil (n = 1). The highest resistance proportion was to streptomycin at 17%, present across all sample types and in combination with resistance to other antimicrobials in 5 of 6 isolates. All isolates were susceptible to amoxicillin/clavulanic acid, nalidixic acid, ampicillin and trimethoprim-sulfamethoxazole, and one chicken isolate showed intermediate susceptibility to ciprofloxacin. For azithromycin, the panel range does not extend to the NARMS resistance breakpoint of ≥ 32 µg/mL. However, all isolates had MICs of ≤ 8 µg/mL, within the tested range and below the susceptible breakpoint of ≤16 µg/mL and were therefore classified as susceptible. Susceptibility to sulfisoxazole could not be determined due to limited drug ranges in this panel and is not reported in this study.

**Table 2.**
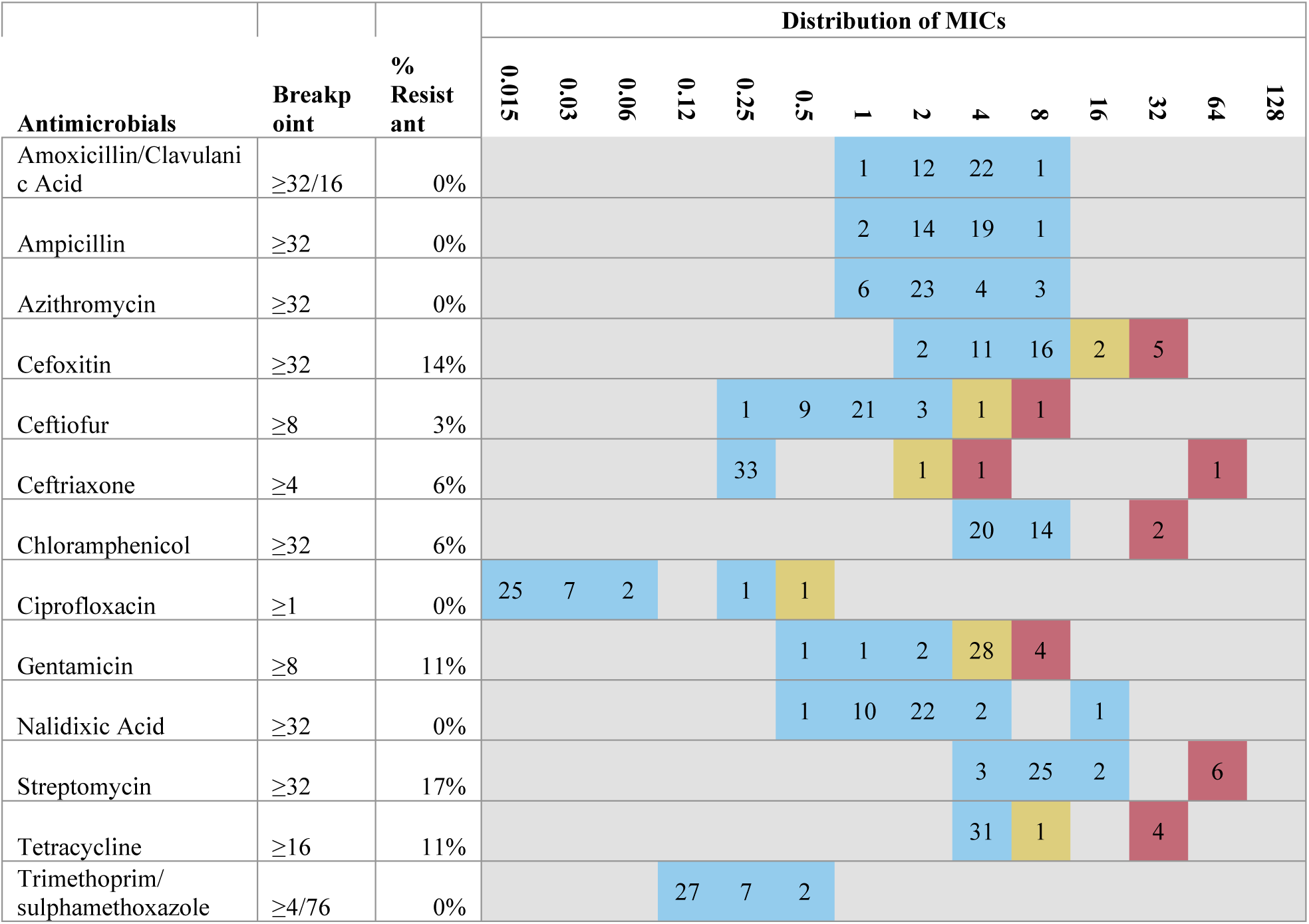
The Minimum Inhibitory Concentration (MIC) distribution and number of *E. coli* isolates at each antimicrobial concentration (n = 36), determined with the Sensititre CMV3AGNF panel. Breakpoints from CLSI were used for all antimicrobials apart from streptomycin and azithromycin breakpoints that were interpreted using NARMS. Blue, yellow and red boxes represent susceptible, intermediate and resistant isolates, respectively. E. coli ATCC 25922 was included as the quality-control organism and results were accepted only when control MICs fell within the CLSI acceptable ranges.

### Whole genome sequencing of isolates

A total of 33 of the 36 *E. coli* isolates with MIC data were sequenced and 31 passed assembly quality control (Supplementary Table S2). Two deer isolates were excluded, one as a mixed culture and one for a fragmented assembly. Short-read assemblies (n = 22) had 43 to 392× coverage, ≥ 99.6% completeness and ≤ 0.94% contamination (4.8 to 5.7 Mbp, 204 to 894 contigs). Nanopore assemblies (n = 9) had 52 to 342× coverage, ≥ 99.9% completeness and ≤ 0.6% contamination (4.8 to 5.9 Mbp, 2 to 16 contigs). The three isolates that were not sequenced (two chicken, one soil) are marked in Supplementary Table S2.

### MLST, serotypes and population structure

As shown in Figure 1, there were 15 different multilocus sequence types (STs) identified across the 31 sequenced strains, and one isolate was not assigned a ST. ST57 was the most common type (n = 6) followed by ST13 (n = 4) and ST7032 (n = 3). The most prevalent serotype was O176:H4 (n = 6), followed by O174:H2 (n = 3) and O146:H21, O91:H14, O124/O164:H25 and O156:H8 (n = 2 each). The largest group was the O176:H4 ST57 isolates (n = 6) from OR1, from sheep feces in both years together with one soil sample from Field B, collected 7 days after the sheep grazing pass. These isolates differed by 1 to 18 core-genome SNPs after recombination masking (Supplementary Table S4 and Supplementary Figure S1), which is consistent with a single lineage circulating on the farm and transferring to soil rather than with independent introductions. Among the six O176:H4 ST57 isolates, the soil isolate AP7001, collected 7 days after grazing in April 2022, differed from the sheep isolate AP6332, collected in July 2021, by one core-genome SNP before and after recombination masking (ANI 100.00%, aligned fraction 99.9%), and the two formed a clade with 96% bootstrap support. The other four sheep isolates differed from AP7001 by 9 to 14 core SNPs after masking. This points to a lineage resident in the flock across both grazing seasons, deposited in soil during the April grazing event and still recoverable seven days later, though the matching genotype was not recovered directly from the April fecal samples. Both isolates in this pair were sequenced on the Nanopore platform and AP7001 served as the mapping reference, whereas three of the four more distant sheep isolates were sequenced on short-read platforms; the sequencing platform of each isolate is given alongside the pairwise distances in Supplementary Table S4. The two ST13 soil isolates (AP7002, AP7749) differed by one core SNP after masking (ANI 99.99%), whereas the two ST13 chicken isolates (AP7710, AP7716) differed from the soil isolates by 868 to 872 core SNPs (ANI 99.57 to 99.61%) which does not support transfer between animals and soil in the ST13 group. AP7002 and AP7749 were recovered from the same plot at OR1 in Field B, a non-productive block grazed continuously by chickens with a single sheep pass, on 19 April and 18 July 2022, the 7 day and 90 day post-grazing timepoints. A single core SNP across that interval indicates that the lineage persisted in the orchard soil over the three months. The chicken isolates sequenced all belonged to a different ST13 lineage, but only a single colony was characterized per sample, so the soil lineage could have been carried by chickens that were not among those sampled, or by a source that was not sampled at all.

**Figure 1:**
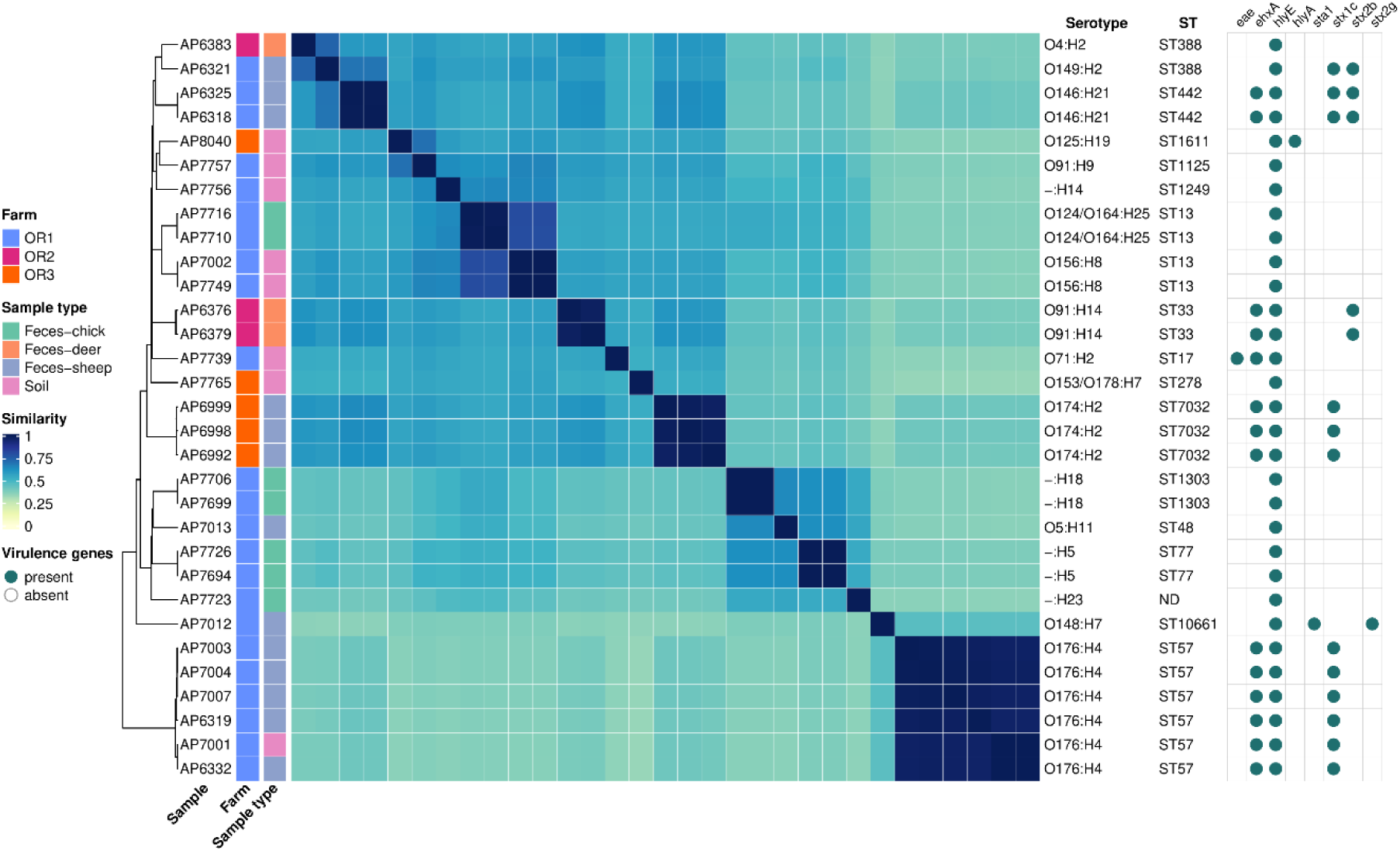
Whole genome similarity, serotypes, sequence types, and virulence gene carriage of *E. coli* isolates (n = 31) from soil and fecal samples from organic nut orchards with grazing animals over 2021 and 2022. Left panel: sourmash plot, pairwise k-mer similarity matrix with k = 31 and scaled value of 10 for each isolate pair. Cell color of white signifies no shared k-mers and dark blue indicating identical isolates. Illumina, DNBSEQ and Nanopore assemblies are shown in the same matrix. Right panel: filled dots mark the pathotype genes and *stx* subtypes detected with ECTyper.

### Virulence factors, AMR and plasmids

From WGS analysis with ECTyper, Shiga toxin genes were detected in 15 isolates: *stx*_1c_ (n = 9), *stx*_1c_;*stx*_2b_ (n = 3), *stx*_2b_ (n = 2) and *stx*_2g_ (n = 1) (Figure 1). The *stx*_1c_ subtype was predominant, present alone or in combination with *stx*_2b_ in 12 of 15 STEC isolates. All O176:H4 and O174:H2 serotype isolates carried *stx*_1c_. Sheep fecal isolate AP7012 carried *stx*_2g_ and the heat-stable enterotoxin *sta1*, making it a STEC/ETEC hybrid. Only one isolate, AP7739, isolated from soil, carried the *eae* intimin gene and Shiga toxin genes were not detected with WGS in this sample. Other virulence genes *ehxA* was detected in 13 of 15 STEC isolates, *astA* in seven isolates (three STEC) and *lpfA* in one *stx*-negative soil isolate, while *saa*, *subAB* and *iha* were not detected (Supplementary Table S3 and S5). Although all isolates were *stx*-positive by colony PCR, 16 of 31 did not have *stx* detection in their assemblies, including all seven chicken isolates and seven of eight soil isolates. Mapping raw reads to *stx* subtype references did not recover *stx* at genome-level depth in any of these 16 isolates (Supplementary Table S3). Repeat PCR was performed on the sequenced isolates, which had been frozen and thawed several times since banking, and on replicate isolates from the same sample bags, which had not been subcultured or freeze-thawed. All seven chicken and six soil sequenced isolates tested were *stx*-negative by PCR. *Stx*-positive replicates were recovered from five bags (three chicken, two soil), showing that *stx*-positive *E. coli* were present and that the original colony PCRs were not false positives; for these, loss of the *stx*-encoding prophage during storage and passage may explain the *stx*-negative genomes. In the remaining eight bags (four chicken, four soil) the unpassaged replicate was also *stx*-negative, so loss of the *stx*-encoding prophage during storage, free *stx-*encoding phage in the enrichment broth or a mixed primary colony are possible explanations.

Screening of AMR genes with CARD/RGI was performed on 31 isolates and 59 unique genes were detected, of which eight were acquired resistance determinants and the remaining 51 were chromosomal genes, efflux and regulatory loci. Only acquired determinants are considered further; the intrinsic loci and complete RGI hits are listed in Supplementary Table S6. For the 31 isolates with matching WGS and MIC data, the four isolates with phenotypic tetracycline resistance AP7001 (soil), AP6332 (sheep feces), and AP7710 and AP7716 (chicken feces), were the only isolates that carried the *tet*(A) gene. Only one isolate, AP7739 isolated from soil, carried the *tet*(C) gene and exhibited intermediate tetracycline resistance. The two isolates with chloramphenicol resistance, AP7001 and AP6332, were also the only isolates with the *floR* gene, demonstrating genotype-phenotype concordance. These isolates also carried the aminoglycoside phosphotransferases genes APH(3’’)-*Ib*, and APH(6)-*Id,* the streptothricin acetyltransferase genes SAT-2, as well as the aminoglycoside nucleotidyltransferase gene ANT(3’’)-*Ia*. Of these, APH(3’’)-*Ib*, APH(6)-*Id* and ANT(3’’)-*Ia* confer the streptomycin resistance that was phenotypically observed, whereas SAT-2 confers streptothricin resistance and does not contribute to the streptomycin phenotype. Cefoxitin resistance (AP6321, AP6383, AP7716), ceftriaxone resistance (AP7716), gentamicin resistance (AP7003, AP7012, AP7749) and streptomycin resistance in AP6383 and AP7716 were not explained by acquired resistance genes, and no chromosomal *amp*C promoter or attenuator mutations or *omp*C/*omp*F truncations were detected in the cefoxitin or ceftriaxone-resistant isolates (Supplementary Table S8).

There were 18 unique plasmid replicons identified across the 31 isolates with an average of two replicons per isolate with a range of 1-5 (Supplementary Table S7). The most prevalent replicon was IncFIB(AP001918), detected in 17 of 31 of isolates across all four sample type categories. Two of the three MDR isolates, AP7001 (soil) and AP6332 (sheep), each carried two replicons IncFIB(AP001918) and IncB/O/K/Z. In both Nanopore assemblies, the IncB/O/K/Z plasmid was a closed, circular 105,665 bp contig carrying all seven acquired AMR genes (*tet*(A), floR, sul2, ANT(3’’)-*Ia*, APH(3’’)-*Ib*, APH(6)-*Id* and SAT-2), consistent with the observed chloramphenicol, tetracycline and streptomycin resistance. The plasmid was identical in length and in acquired gene content in the two assemblies, which were generated independently from different isolates, and this agreement is independent of the sequencing platform and of the single-nucleotide distance between the two chromosomes. The other four ST57 sheep STEC (AP6319, AP7003, AP7004, AP7007) carried only IncFIB(AP001918) and lacked the IncB/O/K/Z plasmid.

## Discussion

While integrating grazing animals has economic and environmental advantages for nut orchards, potential food safety risks have impacted the adoption of this practice, and guidelines restrict grazing timeframes. In this pilot study, spring grazing with sheep and chickens in three nut orchards did not result in persistent environmental pathogen contamination. Specifically, pathogenic organisms that have been linked to nut outbreaks in California, *E. coli* O157:H7 and *Salmonella* (16,57), were not detected in any fecal samples or soil across this study. Additionally, despite STEC fecal shedding prevalence of 34.2% across chicken, sheep and wildlife, evidence of soil contamination with STEC was rare, detected by PCR in only 3.2% of soil samples. This indicates that livestock carriage of pathogens does not consistently result in transfer or persistence of pathogens in the orchard soil. These findings align with recent cover crop and livestock integration studies in vegetable fields reporting low soil STEC prevalence rates of 0.1% and 1.4% following grazing by sheep with STEC fecal shedding (58,59). Sutherland *et al.* (2009) also reported no detection of *E. coli* O157:H7 or *Salmonella* in sheep feces and grazed soil in ten Ontario sheep pastures. Overall, these findings challenge the perception that livestock in crop production environments represent a uniformly higher risk to food safety, instead, risk likely depends on multiple interacting factors including the pathogen virulence, wildlife activity, environmental conditions, animal species, herd-specific pathogen prevalence and on-farm management practices (61).

Pathogen survival in orchard soils was limited, with STEC detections likely attributable to reintroduction rather than environmental persistence following sheep grazing. This was demonstrated in OR1, where an O176:H4 ST57 lineage detected in sheep feces across both years was recovered from soil 7 days after the sheep grazing pass but was absent from all later soil samples. That soil isolate differed by one core-genome SNP from a sheep isolate collected nine months earlier and shared the same closed multidrug resistance plasmid, whereas sheep sampled in the same block on the same day carried isolates that differed from it by 9 to 14 core SNPs. The nine-month interval indicates a lineage resident in the flock across both seasons, and its absence from later soil samples indicates transfer to soil without subsequent persistence. STEC was not detected in any soil samples from two fields despite sheep fecal prevalence of 27.8% at OR1, where Field A and Field B flocks were sampled together, and 15% at OR3. The highest soil contamination prevalence of 9.4% was observed at OR1 Field B in 2022, where chicken feces and soil both tested positive between D7 and D90 before clearing by harvest. Continuous chicken grazing at this site may have reintroduced *E. coli* throughout the sampling period, however, this was not supported by WGS data. These ST13 soil isolates differed from the ST13 chicken isolates by approximately 870 SNPs, and the only STEC from Field B soil confirmed by WGS (AP7001) matched a sheep isolate. Instead, the two ST13 isolates recovered from Field B soil differed by a single core SNP and came from the same plot three months apart, indicating that this lineage persisted in the orchard soil between visits, although *stx* genes were not detected in either assembly. The interpretation of STEC recovered from OR3 Field C soil at D90 and D120+ post grazing is confounded by sheep escaping into the field prior to sampling and proximity of this site to permanent animal housing. These findings contrast with Diaz *et al*. (2022) who reported persistent STEC and *Salmonella* (27–29% and 22–29%, respectively) in Oklahoma pecan orchard soils, two and four months after cattle grazing. This difference may be due to harsh Californian orchard conditions that are characterized by dry soil, heat and UV exposure from widely spaced tree rows that may reduce potential STEC survival. Additionally, sheep produce smaller, faster-drying fecal pellets compared to cattle, and STEC survival rates are known to differ across host fecal matrices (63). Read together with the present data, this indicates that interval length alone is a weak control point and that animal species, the drying rate of the fecal matrix, local climate and continuous introduction from resident animals are the stronger determinants of whether viable pathogens are present in orchard soil at harvest. More longitudinal sampling that quantifies pathogen decline in soil is needed to test this directly.

All STEC isolates identified by WGS in this study were assessed as low public health risk, with none belonging to the ’Big Six’ serogroups (O157, O26, O103, O45, O111, O145) and no detection of the Shiga toxin subtypes most associated with severe human disease (*stx*_2a_, *stx*_2c_, and *stx*_2d_) (64). Following the World Health Organization framework, which categorizes STEC by *stx* subtype and the presence of adhesion factors intimin (*eae)* or the enteroaggregative regulator *aggR* (65), all isolates fell into the lowest level risk category, associated with possible diarrhea but not with hemolytic uremic syndrome (HUS) or bloody diarrhea. The *eae* gene was detected in only one soil isolate (AP7739), with no *eae* detection in animal derived STEC isolates, consistent with the low prevalence of *eae* reported in sheep STEC in China (66) and the Caribbean (67). Shiga toxin subtype *stx_1c_* is considered less pathogenic overall (68) and was the predominant subtype in sheep fecal STEC (11 of 12), comparable to sheep STEC studies in Ireland, Switzerland and China reporting this subtype in 57-66% of isolates (66,69,70). Subtype *Stx*_2b_ was identified in both deer STEC isolates, consistent with previous reports of this subtype in 70% of deer meat STEC (71) and 73% from deer fecal STEC (72). However, *stx* genes detected in initial PCR were not identified in WGS in 16 of 31 isolates, including all chicken fecal isolates, impeding meaningful analysis of the virulence potential of these isolates. On repeat PCR of replicate isolates banked from the original sample bags, which had not been subcultured or freeze-thawed, *stx*-positive isolates were recovered from five samples (three chicken, two soil), indicating that *stx*-positive *E. coli* were present and the original PCRs were likely not false positives. This discordance has been reported previously as the loss of Shiga toxin genes is consistent with known instability of *Stx*-encoding prophage *in vitro*, and loss of these genes during lab passaging and storage is an issue reported in other STEC studies (69,73–75).

Surveillance of AMR in STEC isolates from integrated crop-livestock systems is critical from a One Health perspective, as antimicrobial use in livestock management and crop production may drive resistance gene selection and impact the future effectiveness of antimicrobials. The overall AMR prevalence was 33% across all sample types, with multidrug resistance observed in 8.3% of isolates, all from OR1. The absence of AMR in OR3, a farm that owns and manages a flock of fully organic sheep, compared to OR1, that rented conventionally managed sheep, indicates that farm-level management practices may influence AMR load. While comparable integrated animal-orchard STEC AMR data are limited, the overall sheep resistance rates of 31% found in this study were lower than the 83% reported in a sheep study in Brazil (76) and 47% in sheep in China (77). On the other hand, a study in Spain (78) reported resistance in 20% of O157:H7 isolates from sheep, while all non-O157 STEC isolates were susceptible. A key finding of this study is that STEC isolates were susceptible to critically important human antimicrobials including fluoroquinolones, ampicillin and trimethoprim-sulfamethoxazole although two isolates were resistant to the third-generation cephalosporin ceftriaxone. Additionally, high-concern acquired resistance genes including the mobile colistin resistance gene *mcr-1* and extended-spectrum β-lactamase enzymes from the *bla*_CTX-M_ family were not detected (79–82). Phenotypic resistance was highest to streptomycin at 17% and was detected in all sample types. This is consistent with the use of this antimicrobial in conventional agriculture as a crop pesticide and historically in veterinary medicine (83) which may contribute to selective pressure in ICLS. Streptomycin is not included in the list of drugs prohibited for extralabel drug use in food-producing animals in the US (84). However, because of the prolonged elimination and residue concern, the American Veterinary Medicine Association does not support use of extra label aminoglycosides (except for neomycin sulfate) in small ruminants. For chicken, streptomycin is approved for use in drinking water as the treatment of nonspecific infectious enteritis under CFR §520.2158, but not for chickens producing eggs for human consumption. The streptomycin resistance rate in this study is lower than the 58.7% rate reported in livestock *E. coli* in the US between 1950-2002 (85) and the 88% streptomycin resistance rate in STEC from farms with co-located animal and produce operations in Tennessee and North Carolina (86). There was an 11% resistance prevalence to gentamicin in the STEC isolates from this study. A higher gentamicin resistance prevalence of 63.6% was previously reported in *E. coli* and other bacterial isolates from sheep and goat fecal samples on smallholder farms in Tennessee and Georgia, where concerns about gentamicin overuse were emphasized (87). The sulfonamide resistance gene *sul2* was detected in two isolates (AP7001, AP6332), on the shared IncB/O/K/Z plasmid. This finding contrasts with the 59% *sul2* prevalence and 47% phenotypic resistance to this drug reported in sheep STEC from Bangladesh (88). Overall tetracycline resistance was 11% in this study, with genotype-phenotype concordance for the efflux pump gene *tet*(A) detected in isolates from OR1, the only farm incorporating chicken grazing. This observation is consistent with the previously reported association of *tet*(A) and poultry-associated *E. coli* (82). According to a National Antimicrobial Resistance Monitoring System study, tetracycline resistance was the most prevalent in generic *E. coli* isolated from sheep, goat, and lamb cecal content samples (89). They reported that this may be linked to the dissemination of *tet* resistance genes likely introduced by the use of tetracyclines in veterinary and agricultural settings. Collectively, the AMR profiles observed in this study reflect a low resistance burden in STEC from integrated crop-livestock systems, with limited resistance detected to antimicrobials of critical clinical importance.

The findings of this study are limited by small farm numbers with heterogeneous grazing animals and practices, and non-standardized sampling timeframes which limits the generalizability of the results. OR1 Field B had continuous chicken grazing, sheep samples were not collected from OR2 and OR3 Field C had sheep escape prior to sampling, making the interpretation of temporal effects difficult for these sites. In addition, *stx* was lost from some isolates between isolation and sequencing, so the virulence profiles of *stx*-negative isolates, including all chicken isolates, could not be assessed. Isolates were also sequenced on three platforms, and SNP distances within the ST57 cluster were calculated against a Nanopore reference, so small distances between Nanopore-sequenced pairs may be favored relative to comparisons that cross platforms; the shared closed plasmid provides corroboration that does not depend on platform. The design is further limited by enrichment-based presence and absence detection without quantification of pathogen or indicator organism concentration, so rates of decline could not be estimated, and by the absence of nut sampling, which means the exposure pathway of interest was inferred from soil rather than measured on the crop. Despite these limitations, the low virulence and AMR profiles observed, combined with transient soil contamination, support the viability of rotational grazing in organic orchard systems from a food safety perspective.

The virulence potential and AMR profiles of STEC isolated from sheep, chicken, wildlife and soil samples across three Californian nut orchards incorporating rotational grazing was assessed in this study. *Salmonella* and *E. coli* O157 were not detected in any sample and the STEC confirmed by sequencing lacked the Shiga toxin subtypes and virulence genes most associated with human disease. Additionally, a low AMR prevalence was reported and there was limited resistance to critically important antimicrobials collectively suggesting a low public health risk from these systems. As California almonds are subject to a mandatory validated treatment achieving a 4-log *Salmonella* reduction before sale, whereas walnuts carry no equivalent requirement, this conclusion is better supported for the almond orchards than for the walnut orchard. Soil contamination was rare and transient, and pathogen deactivation was mostly observed to fall within grazing to harvest temporal guidelines.

## Methods

### Study design

The sampling design consisted of a repeated cross-sectional study of three integrated crop-livestock farms over two growing seasons during 2021 and 2022. Farms were selected across three regions of California, including the Sacramento Valley, San Joaquin Valley, and Capay Valley. The inclusion criteria for farm selection were: 1) the farm incorporates crop-livestock and rotational grazing systems with poultry or small ruminants; 2) the size of the farm is small to medium (<500 goats/sheep; < 1,000 chickens/year), and uses direct-marketing; 3) the farm contains established tree nut orchards and is organically managed and certified; 4) the farms have established grazing practices with sheep and/or chickens incorporation over two to four years; 5) willingness to participate. Two almond and one walnut orchard with sheep and/or chicken grazing were enrolled in the study. Each farm was visited once prior to grazing and then at least once post grazing with 1-2 visits conducted depending on the grazing schedule for each orchard during the respective growing season. Sampling timepoints are reported as days after the most recent grazing pass (D0: baseline or taken prior to grazing, D7: 7 days post-grazing). D120+ denotes samples collected 120 or more days after the last grazing pass. The walnut farm (OR1) had two fields, Field A with sheep grazing only (0.15 km²), and Field B (0.38 km²), a non-productive orchard with semi-permanent chicken and sheep grazing. OR1 rented their grazing animals and these were conventionally managed. The almond farm OR2 also had two fields, Field A (0.03 km²) and Field B (0.02 km²) that were both grazed with rented, conventionally managed sheep, the spring prior to the study start only. Due to farmer constraints, these fields were only grazed once but were included in this study to observe long term pathogen dynamics in grazed orchards. Farm 3 (OR3) was also an almond farm with four fields and sheep grazing only. This farm owned their sheep and these were managed with organic practices. OR3 Field A (0.27 km²) was included as a control site with no grazing passes. Field D (0.38 km²) and Field B (0.11 km²) both had one grazing pass with sheep. Field C (0.47 km²), located directly next to the sheep barn, was grazed more frequently, with four grazing passes.

### Sample collection methods

For fecal sample collection of each flock, 8 to 10 individual fresh sheep fecal deposits of approximately 30 g were collected using sterile scoops from the ground and placed into sterile Whirl-Pak bags (Nasco, Modesto, California, USA). For chicken fecal samples (OR1 only), 10 g of composite fecal samples consisting of three individual fecal droppings for each coop were also collected. Wildlife (deer) fecal samples were opportunistically collected when observed in the orchard soil during visits (OR2 only). In total, 79 fecal samples were collected, 56 from sheep, 15 from chicken fecal and eight from wildlife/deer.

For soil sample collection in each field, eight soil samples consisting of five composite soil cores collected in a W shaped pattern in the space of five by five trees were collected (28). Composite soil cores were collected with a 2.5 cm diameter metal soil coring tool to a depth of 15 cm and placed into sterile Whirl-Pak bags before massaging to homogenize soil cores. Sampling grids were randomly selected using satellite images of the nut orchards and a buffer of three trees was used at each site to control for any edge effects. A total of 284 soil samples before and up to 120 days post-grazing were collected in total.

### Pathogen isolation methods

Previously described culturing methods were used to detect for the presence of foodborne disease pathogens *E. coli* O157:H7 (29), STEC (29,30) and *Salmonella enterica* (31). Beginning with a generic enrichment, 10 g of fecal samples or 30 g of soil samples were placed in 90 mL or 270 mL of tryptic soy broth (TSB, Becton Dickinson, Sparks, Maryland, USA) respectively. Samples were massaged manually for 1 min to homogenize before incubation with orbital shaking at 100 rpm for 2 h at 25 °C, then 8 h at 42 °C and holding with no agitation at 6 °C in a Multitron shaking incubator (Eppendorf, Hauppauge, New York, USA).

Immunomagnetic separation (IMS) was performed on the primary TSB enrichment broth to isolate *E. coli* O157:H7 with Dynabeads™ anti-E*. coli* O157 (Thermo Fisher Scientific, Vilnius, Lithuania) and the automated Dyna BeadRetriever™ (Invitrogen, Carlsbad, California, USA). The elution was then streaked onto Sorbitol MacConkey II Agar with Cefixime and Tellurite (CT-SMAC, Becton Dickinson, Sparks, Maryland, USA) with presumptive positive isolates appearing colorless after incubation for 24 h at 37 °C. The IMS elution was also streaked in parallel onto Rainbow Agar O157 (Biolog, Hayward, California, USA) containing novobiocin (20 mg/L, Sigma-Aldrich, St. Louis, Missouri, USA) and potassium tellurite (0.8 mg/L, Sigma-Aldrich, St. Louis, Missouri, USA) and incubated for 24 h at 37 °C to detect presumptive isolates with black or grey colony morphologies. Isolates were then streaked onto the alternative media for two passages before streaking on Tryptic Soy Agar (TSA, Becton Dickinson, Sparks, Maryland, USA) with incubation for 24 h at 37 °C and confirmation with PCR for the detection of the *E. coli* O157:H7 *eaeA* gene (32).

For STEC enrichment and isolation, 1 mL of the primary TSB enrichment broth was added to 9 mL of modified enterohemorrhagic E. coli (mEHEC) selective broth (Biocontrol, Bellevue, Washington, USA) and incubated at 42 °C for 12 h. This enrichment broth was then streaked onto ChromSTEC (CHROMagar™, Saint-Denis, France) and incubated at 37 °C for 24 h where presumptive STEC colonies, indicated by mauve or purple color and fluorescence under ultraviolet light, were isolated and further purified with two passages on ChromSTEC. Up to six STEC isolates were collected from each sample bag. Presumptive STEC colonies were then streaked on TSA with incubation for 24 h at 37 °C. STEC isolates were confirmed with PCR assays to detect for presence of *stx* _1_ or *stx* _2_ genes (33).

For *Salmonella* isolation, 1 mL of the primary TSB enrichment was aliquoted into 9 mL of Buffered Peptone Water (BPW, Hardy Diagnostics, Santa Maria, CA, USA) before incubation for 24 h at 37 °C. From this solution, 1 mL was added to 10 mL of Tetrathionate (TT) broth and 0.1 mL was added to 10 mL of Rappaport-Vassiliadis (RV) broth (BD, Sparks, MD) with both tubes incubated at 42 °C for 24 h. Enrichments were plated onto Xylose Lysine Desoxycholate Tergatol-4 (XLT4) agar (Becton Dickinson, Le Pont de Claix, France) 37 °C for 24 h with presumptive positive colonies with black colony morphologies purified with up to two passages on XLT4 before streaking on TSA with incubation for 24 h at 37 °C and confirmation with PCR through detection of the *invA* gene (32). All confirmed pathogens were stored at -80 °C in 15 % glycerol and TSB and resuscitated on TSA overnight incubation at 37 °C for subsequent testing.

### Antimicrobial susceptibility testing

For isolated pathogens, the determination of minimum inhibitory concentrations (MICs) of veterinary related antibiotics was performed using standard broth microdilution methods (Sensititre™ CMV3AGNF, ThermoFisher, West Sussex, United Kingdom) and interpreted using breakpoints from Clinical and Laboratory Standards Institute (CLSI) M100 (34) for Amoxicillin/Clavulanic Acid, Ampicillin, Ceftiofur, Cefoxitin, Ceftriaxone, Chloramphenicol, Ciprofloxacin, Gentamicin, Nalidixic Acid, Tetracycline, and Trimethoprim/sulphamethoxazole. As CLSI breakpoints for Streptomycin and Azithromycin are not available for *E. coli*, the National Antimicrobial Resistance Monitoring System (NARMS) breakpoints were used for interpretation of these MICs (35). Susceptibility to Sulfisoxazole could not be determined due to limited drug ranges in this panel and is not reported in this study. Multidrug resistant isolates were determined by phenotypic resistance to at least three antimicrobial classes. *E. coli* ATCC 25922 was included as a quality-control organism and results were accepted only when control MICs fell within the CLSI acceptable range. Breakpoints and QC results are listed in the Supplementary data Table S1.

### Whole genome sequencing

One isolate from each soil or fecal sample with pathogen detection was prepared for whole genome sequencing (WGS). DNA extraction and quality assessment were completed following established methods (36,37). Briefly, DNA was extracted using the Wizard genomic DNA purification kit (Promega Corporation, Madison, WI, USA), according to the manufacturer’s instructions for Gram-negative bacteria. DNA purity and yield were measured with the NanoDrop One UV-Vis Spectrophotometer (Thermo Scientific, Waltham, MA, USA) with quality thresholds set at A260/280 ≥ 1.8 and A260/A230 ≥ 2.0. The integrity of the DNA was analyzed with the TapeStation System 4200 with the Genomic DNA ScreenTape Assay (Agilent, Santa Clara, CA, USA). Library preparation and sequencing were completed by the laboratory of Dr. Bart Weimer (UC Davis) with the 100K Pathogen Genome Project methods (38,39). Libraries were constructed using the KAPA HyperPlus Library Preparation Kit (Roche, Indianapolis, IN, USA) with 450 bp average inserts and sequenced on the Illumina HiSeq X Ten with PE150 (paired-end sequencing with a read length of 150 bp) (Illumina Inc., San Diego, CA, USA). Isolates AP6318, AP6319, AP6321, AP6325 and AP7004 were sequenced on a DNBSEQ-G400 (MGI Tech) with 150 bp paired-end reads. For the short reads, low-quality sequences and adapters were removed with Trimmomatic (v 0.39) (40), FastQC was used to check the sequence quality (v 0.12.1) (41) followed by assembly using Shovill (v 1.1.0) with the SPAdes assembler (42). Mosdepth (v 0.3.6) (43) was used to assess coverage and depth. The nine isolates (AP6332, AP7001, AP7002, AP7003, AP7706, AP7726, AP6376, AP7765 and AP7013) were sequenced using the Rapid Barcoding Kit 94 V14 (SQK-RBK114.96) for library construction and loaded onto R10.4.1 flow cells (FLO-MIN114) for sequencing on the GridION (Oxford Nanopore Technologies) following the manufacturer’s instructions (44). Of 26 isolates sequenced on short-read platforms, two mixed cultures were resequenced on Nanopore and two were excluded, one as a mixed culture and one for a highly fragmented assembly. A further seven isolates were sequenced on Nanopore only, giving 31 isolates for analysis (Table S2). Nanopore sequences were processed with base-call model dna_r10.4.1_e8.2_400bps_hac@v4.3.0 and assembled using the EPI2ME pipeline (wf-bacterial-genomes v1.4.2) with Flye (v 2.9.6) (45). Completeness (≥99%) and contamination (≤1%) were assessed for all assemblies with CheckM (v 1.2.2) using the workflow ’lineage_wf’ (46). Bracken with Kraken2 (47) was used to assess contamination and species identity.

Multilocus sequence typing (MLST) was performed using MLST (v 2.35) with the *ecoli_achtman_4* scheme (48). Antimicrobial resistance genes were identified using Resistance Gene Identifier (RGI) (v 6.0.8) against the Comprehensive Antibiotic Resistance Database (CARD) (49) with strict and perfect cutoffs and ≥90% identity. Chromosomal *amp*C promoter and attenuator mutations and porin (*omp*C, *omp*F) mutations were screened with AMRFinderPlus (v4.2.7, downloaded 7 Aug 2026) (50). ABRicate (v 1.3.1) (51) was used to screen contigs for virulence factors against the ecoli_vf database (52) (downloaded 5 March 2026) with thresholds of ≥90% coverage and ≥95% identity. In silico serotyping and shiga toxin subtypes, intimin (*eae*) and enterohemolysin (*ehx*A) were called in silico with ECTyper (v 2.0.0) (53). Plasmids were identified using PlasmidFinder (v3.0.0, database v2.2.0) against the Enterobacteriales database retaining hits with ≥90% coverage and ≥95% identity (54).

To distinguish loss of *stx* from collapse of the *stx*-encoding prophage during assembly, raw reads of all isolates were mapped with minimap2 (v2.28) to 151 *stx*1 and *stx*2 subtype alleles from the ECTyper database (v2024-10-03) and read depth across the best-matching allele was calculated with samtools (v1.20) and compared with genome-wide coverage (Table S3). Reads aligned over ≥50 bp at ≥90% identity were counted. Repeat *stx* PCR, with replicates, was performed on less-passaged original glycerol stocks of all banked isolates from the chicken samples and from selected soil samples.

To compare genetic similarity of isolates, Sourmash (version 4.9.4) (55) was used with a k-mer size of 31, a scaled value of 10 and the Jaccard similarity metric, with pairwise comparisons plotted in an all-by-all heatmap. As k-mer Jaccard similarity is sensitive to assembly fragmentation and platform-specific error, candidate transmission clusters, defined as sequence types containing both a soil and an animal fecal isolate (ST57 and ST13), were compared by average nucleotide identity (skani v0.3.2) and by reference-based single-nucleotide polymorphism (SNP) calling. Within each cluster, assemblies were aligned with Snippy (v4.6.0) to the assembly of the cluster’s soil isolate (AP7001 for ST57, AP7002 for ST13). A core-genome alignment was built with snippy-core and pairwise core SNP distances were calculated with snp-dists (v1.2.0). Recombinant regions were removed with Gubbins (v3.4.3). Post-masking distances and trees were computed from the masked alignment over sites called in every isolate, with invariant sites retained. Trees were generated with IQ-TREE (v2.0.7) with a GTR+G model and 1000 ultrafast bootstrap replicates.

## Statistical analysis

Data analysis was conducted using R (v 4.1.2). Descriptive statistics were used to examine the numbers of pathogen positive samples, prevalence of AMR and VF genes and plasmids. Prevalence over the overall two-year study period was reported with Wilson 95% confidence intervals, whereas Clopper-Pearson exact 95% confidence intervals were calculated for each field at each sampling timepoint because of the small sample sizes. Because 8, 12, 16, or 24 samples were collected per field at each timepoint, the upper 95% confidence bound for field-timepoints with no detections was 36.9%, 26.5%, 20.6%, and 14.2%, respectively. These detection limits are reported alongside all negative results. MIC data was presented in squashtograms. Heatmaps were generated using the ComplexHeatmap package in R (56).

## Data Availability

The datasets generated and analyzed during the current study are available in the National Center for Biotechnology Information (NCBI) repository under BioProject accession number PRJNA1555146. The raw sequencing reads have been deposited in the Sequence Read Archive (SRA) and the accession numbers are listed in the Supplementary Table S2.

## Acknowledgements

This work was supported by the USDA National Institute of Food and Agriculture, Center for Food Animal Health, Animal Health (Project No. CALV-AH-406). We thank the participating farmers for access to their orchards, Sara Tiffany for outreach and for facilitating participant enrollment, and Carolyn Chandler-Khayd, Thomas Luder, Nicole Tng, Nicholas La and Olivia Thorn for assistance with sampling and laboratory work.

